# ABCB1 overexpression through locus amplification represents an actionable target to combat paclitaxel resistance in pancreatic cancer cells

**DOI:** 10.1101/2023.05.30.542412

**Authors:** Cecilia Bergonzini, Alessandro Gregori, Tessa M.S. Hagens, Vera E. van der Noord, Bob van de Water, Annelien J.M. Zweemer, Mjriam Capula, Giulia Mantini, Asia Botto, Francesco Finamore, Ingrid Garajova, Liam A. McDonnell, Thomas Schmidt, Elisa Giovannetti, Erik H.J. Danen

## Abstract

**Aims:** Chemotherapies such as gemcitabine/nab-paclitaxel are confronted with intrinsic or acquired resistance in pancreatic ductal adenocarcinoma (PDAC). We aimed to identify novel actionable mechanisms to overcome such resistance.

**Methods:** Three paclitaxel (PR) and gemcitabine resistant (GR) PDAC models were established. Transcriptomics and proteomics were used to identify conserved mechanisms of drug resistance. Genetic and pharmacological approaches were used to overcome paclitaxel resistance.

**Results:** Upregulation of ABCB1 through locus amplification was identified as a conserved feature unique to PR cells. ABCB1 was not affected in any of the GR models and no cross resistance was observed. The ABCB1 inhibitor verapamil or siRNA mediated ABCB1 depletion sensitized PR cells to paclitaxel and prevented efflux of ABCB1 substrates in all models. ABCB1 expression was detected in PDAC patients that had received gemcitabine/nab-paclitaxel treatment. A pharmacological screen identified known and novel kinase inhibitors that attenuate efflux of ABCB1 substrates and sensitize PR PDAC cells to paclitaxel.

**Conclusion:** Upregulation of ABCB1 through locus amplification represents a novel, conserved mechanism of PDAC paclitaxel resistance. ABCB1 has not been previously implicated in PR PDAC. The synthetic lethal interactions identified in this study can be further (pre)clinically explored as therapeutic strategies to overcome paclitaxel resistance in PDAC.

## 1. INTRODUCTION

Pancreatic ductal adenocarcinoma (PDAC) is one of the most lethal cancers worldwide (Sung et al. 2021), with a 5-year overall-survival reached in only 10% of patients(Miller et al. 2022). This poor prognosis is due to a lack of early biomarkers, limited therapeutic options, and inherent or acquired chemoresistance (Sarantis et al. 2020; Puik et al. 2022). Currently, surgical resection is the only curative option for patients diagnosed with PDAC but <20% are diagnosed at an early stage and therefore eligible for surgery. In all other cases (*i.e*., advanced PDAC), chemotherapy using FOLFIRINOX or gemcitabine plus nab-paclitaxel are the only treatment options (Milella et al. 2022; Robatel and Schenk 2022). Unfortunately, these chemotherapy regimens increase survival up to 13 months at most, largely due to development of chemoresistance. So far, immunotherapy has not been successful for PDAC patients and new therapies under investigation targeting the tumor or the tumor microenvironment have not reached the clinic (Ayasun et al. 2022; Chakrabarti et al. 2022; Turpin et al. 2022). Therefore, identifying strategies to combat PDAC resistance to currently used chemotherapies is of crucial importance.

Gemcitabine is a cytotoxic DNA-intercalating drug, which arrests aberrant cell proliferation. Chemoresistance to gemcitabine in PDAC has been extensively studied and reported to be multifactorial (Grasso et al. 2017). On the other hand, paclitaxel is a microtubule-stabilizing drug that impedes cell division leading to replication errors and cell death. Paclitaxel also potentiates gemcitabine efficacy by increasing intratumor uptake and inhibiting its inactivation by catabolizing enzymes (Frese et al. 2012; Grasso et al. 2017). Mechanisms underlying paclitaxel resistance in PDAC are poorly understood (Comandatore et al. 2022; Marin et al. 2022). Three studies have investigated paclitaxel resistance in PDAC, reporting that it involves metabolic adaptation (Braun et al. 2020), sustained c-MYC activation (Parasido et al. 2019), and expression of orexin receptor type 1 (Voisin et al. 2022).

ATP-binding cassette (ABC) transporters are responsible for active transport of many substrates, including cytotoxic drugs, across the cell membrane towards the extracellular space. ABC transporters are therefore known as multidrug resistance pumps (Vasiliou et al. 2008; Marin et al. 2022). The ABC family consists of 49 members, among which ABCB1 (also known as MDR1 or P-glycoprotein, P-gp) has been extensively studied in cancer. Increased expression of ABC transporters has been associated with PDAC resistance to gemcitabine and simultaneous downregulation of multiple ABC transporters, including ABCB1, can sensitize PDAC cells to gemcitabine (Chen et al. 2012; Song et al. 2013; Cao et al. 2015; Lu et al. 2019; Marin et al. 2022). Paclitaxel also represents an ABCB1 substrate and ABCB1 has been shown to mediate paclitaxel resistance in several cancer types (Jaramillo et al. 2018) but this has not been addressed in PDAC.

In the present study, we have generated three independent paclitaxel- and gemcitabine resistant PDAC models and found that ABCB1 is amplified in paclitaxel resistant, but not in gemcitabine resistant PDAC cells. We show that pharmaceutical or genetic inhibition of ABCB1 effectively restores paclitaxel sensitivity in the resistant cell lines. We show that ABCB1 is expressed in PDAC patients that had received gemcitabine/nab-paclitaxel treatment. Moreover, as clinical trials of currently available ABCB1 inhibitors have not proven successful due to lack of efficacy or toxicity (Szakács et al. 2006; Jaramillo et al. 2018), we screened a kinase-inhibitor (KI) library for KIs that attenuate efflux of ABCB1 substrates, thereby overcoming paclitaxel resistance. We identified several novel KIs that can be further (pre)clinically explored as therapeutic strategies in combination with paclitaxel to overcome paclitaxel resistance in PDAC and other cancers.

## 2. METHODS

### 2.1 Materials and cell culture

Three PDAC cell lines were used: Patu-T (mesenchymal phenotype), kindly provided by Dr. Irma van Die (Amsterdam UMC, Amsterdam, The Netherland), Suit-2.028 (epithelial phenotype) and Suit-2.007 (mesenchymal phenotype), kindly provided by Dr. Adam Frampton (Imperial College London, London, UK). Patu-T were maintained in DMEM, supplemented with 10% heat-inactivated bovine fetal serum and 1% penicillin/streptomycin, while both Suit-2 cell lines were cultured in RPMI supplemented as described above. All cells were kept in humidified atmosphere of 5% CO_2_ and 95% air at 37 °C, subcultured twice a week, tested monthly for mycoplasma contamination by MycoAlert Mycoplasma Detection Kit (Westburg, Leusden, The Netherlands) and cell identity was verified by short tandem repeats (STR) profiling.

Gemcitabine was kindly provided by Eli Lilly Corporation (Indianapolis, IN, USA) and dissolved in sterile water. Paclitaxel and verapamil were obtained from Sigma (T7402 and V4629, Sigma-Aldrich, St. Luis, MO, USA).

### 2.2 Generation of resistant cell lines

To establish gemcitabine-resistant (GR) and paclitaxel-resistant (PR) cell lines, concentrations causing 50% reduction in cell growth (IC_50_) were determined in parental cells. Cells were then exposed to the respective IC_50_ of the drug and grown for at least 2 weeks with the drug until reaching 80% confluency. After acquiring resistance, the drug concentration was doubled (2x IC_50_) and cells were cultured until they could grow to confluence. The process was repeated with stepwise increasing drug concentrations until the maximum tolerated concentration was reached after 6-12 months. Parental cells never exposed to the drug were cultured in parallel with the resistant cells. To determine stable resistance PR and GR cells were grown in drug-free medium and baseline growth and resistance to the maximum tolerated concentration was analyzed by SRB assay at regular intervals for up to 2 months. The resistance factor was calculated as the ratio of the IC_50_ of resistant versus IC_50_ of parental cells. For IC_50_ > 12 µM, the resistance factor was calculated using the maximum drug concentration used in the SRB assay. Batches of resistant cells used in experiments were maintained in drug-free medium ≤ 2 months.

### 2.3 Immunohistochemistry

Expression of ABCB1 in PDAC patients was evaluated by immunohistochemistry (IHC) in paraffin-embedded tumor specimens from 24 PDAC patients who had been treated with gemcitabine (1,000 mg/m^2^) plus nab-paclitaxel (125 mg/m^2^) in the adjuvant or metastatic (after tumor relapse) setting. All specimens were obtained after patient’s written consent approved by the Ethics Committee of “Area Vasta Emilia Nord” (protocol code 12003 - 17/03/2021). Tissue sections were stained overnight with rabbit anti-human ABCB1 (E1Y7S, mAb #13978; Cell Signaling Technology; dilution 1:400). Sections were reviewed independently by two researchers blinded to clinical data, who scored the immunostaining on the basis of staining intensities and number of stained cells as “low” or “high”. Overall survival (OS) was calculated from the date of pathologic diagnosis (i.e., the date of surgery/biopsy) to the date of death. OS curves were constructed using Kaplan-Meier method, and differences were analyzed using log-rank test with SPSS v.25 statistical software (IBM).

### 2.4 SRB assay

For sulforhodamine B (SRB) assay, cells were seeded in 96-well flat bottom plates at a density of 3000-4000 cells/well. After 72h of drug exposure, plates were fixed with 50% TCA, and incubated with 0.4% SRB at room temperature avoiding light. Plates were washed with 1% acetic acid to remove unbound SRB and air dried. 10 mM Tris was used to extract protein-bound dye and optical density was measured with a BioTek Synergy HT plate reader (SN 269140, BioTek Instruments Inc.) at 490 and 540 nm. IC_50_ was determined through interpolation in GraphpadPrism (version 9.0, Intuitive Software for Science, USA).

### 2.5 RT-qPCR and DNA-qPCR

For RT-qPCR, RNA was isolated with an RNEasy Plus Mini kit (QIAGEN, Cat. 74136). 800 ng RNA was used to generate cDNA with the Thermo Scientific RevertAid H Minus First Strand cDNA Synthesis Kit (Thermo Fisher Scientific, Waltham, MA, USA). For DNA-qPCR, genomic DNA was isolated with the GenElute™Mammalian Genomic DNA Miniprep kit, following manufacturer instructions (Cat. GIN350, Sigma-Aldrich, St. Luis, MO, USA). RNAse A solution (provided with the kit, 1:10 dilution) was used to obtain RNA-free DNA. 7.5 ng of genomic DNA was used as template. qPCR was performed in triplicate using the PowerUp™ SYBR™ Green Master Mix (Thermo Fisher Scientific, Waltham, MA, USA) in a QuantStudioTM 6 Flex Real-Time PCR system (Applied Biosystems®, ThermoFisher Scientific, Waltham, MA, USA). Primers were from Sigma-Aldrich (St. Louis, MO, USA) and were directed against exon-exon boundaries for RT-qPCR or directed against intron-exon boundaries or introns for DNA-qPCR (Table S1). Relative mRNA expression and relative DNA amount was calculated using the 2^-(ΔΔCt)^ method with *ACTB* and *GAPDH* as reference genes.

### 2.6 Western blot

Cells were lysed with RIPA buffer supplemented with 1% protease/phosphatase inhibitor cocktail (PIC, Sigma-Aldrich, St. Luis, MO, USA), 40 µg lysates were separated by SDS–polyacrylamide gel electrophoresis and transferred to PVDF membranes. Membranes were incubated overnight at 4°C with rabbit-anti-human ABCB1 (E1Y7B; mAb #13342; Cell signaling Technology; dilution 1:1000) and mouse-anti-human β-actin (sc-47778; Santa Cruz Biotechnology, Santa Cruz, CA, USA; dilution 1:1000) antibodies, followed by incubation with HRP-conjugated anti-rabbit (#7074; Cell Signaling Technology; dilution 1:2000) and Alexa Fluor® 647-conjugated anti-mouse (115-605-146; Jackson ImmunoResearch, Bio-Connect, Huissen, The Netherlands; dilution 1:1000) secondary antibodies for 1 hour at room temperature. Signals were detected using enhanced chemiluminescence (ECL) and fluorescence readouts.

### 2.7 Hoechst exclusion assay

Cells were seeded at a density of 7000 cells/well in 96-well flat bottom black imaging plates (#655090; Greiner Bio-One™). Cells were allowed to attach for 8 hours and then treated with 1 µM of the selected KI, 10 µM verapamil, or DMSO as negative control. Each condition was tested in triplicate wells. 24 hours after seeding, 1 µg/mL Hoechst33342 was added followed by an additional 2-hour incubation. Subsequently, all wells were aspirated and received fresh medium including the respective inhibitors, without Hoechst33342 and plates were placed in a Nikon TEi2000 confocal microscope equipped with an automated stage and temperature and CO_2_-controlled incubator for live imaging. Total intensity of nuclear Hoechst33342 was calculated with CellProfiler (Carpenter et al. 2006) after watershed segmentation in FIJI-ImageJ (Schindelin et al. 2012). The intensity values of three images from three replicate wells were averaged for each condition. Values of experimental groups were normalized to those of the DMSO control group.

### 2.8 siRNA mediated ABCB1 and Sorcin knockdown

Cells were reverse transfected with 50 nM SMARTpool siGENOME siRNAs (Dharmacon) using INTERFERin transfection reagent (Polyplus; 409-50). A mixture of siRNAs targeting all kinases in the human genome, diluted to a total concentration of 50 nM with a concentration for each individual siRNA ∼0.05 nM was used as control (siKINASEpool). Medium was refreshed after 24 hours. For SRB assays, 3000 cells/well were seeded in triplicate wells in 96-well flat bottom plates. For RT-qPCR, 120,000 cells/well were seeded in duplicate wells in 24-well plates. At 48h and 72h post-transfection, cells in 24-well plates were processed for RT-qPCR and cells in 96-well plates were incubated for an additional 72 hours in presence of DMSO or paclitaxel and subsequently processed for SRB assay.

### 2.9 Extrachromosomal DNA analysis

Cells in the exponential growth phase (70% confluent) were treated with colcemid (KaryoMax, #15212012, Gibco™) for 1-2h at a final concentration of 0.1 µg/ml. Cells were then detached by trypsinization, collected, and treated with a hypotonic solution (75 mM KCl) for 15 minutes. Next, cells were fixed in Carnoy’s fixative (3:1 Methanol:Glacial acetic acid), washed three times and resuspended in 200 μL of Carnoy’s solution. Finally, metaphase chromosomes were prepared by dropping the cell suspension onto glass slides and mounted with ProLong™ Diamond Antifade mountant containing DAPI (Invitrogen, P36966, Waltham, MA, USA). Chromosomes and extrachromosomal DNA (ecDNA) were visualized with a Nikon TEi2000 confocal microscope, with a 60X objective and a 2x digital magnification.

### 2.10 Kinase inhibitor screening

The L1200 library from Selleckchem® (Munich, Germany) was used, containing 760 KIs that were dissolved in DMSO or water at a concentration of 10 mM or 1 mM. 3000 cells/well were seeded in 96-well flat bottom plates. After 24h cells were treated with DMSO only (0.1%), 1 µM KI and DMSO, or 1 µM KI in combination with 0.1 µM paclitaxel. 10 µM verapamil was used as a positive control. After 72 hours, cells were fixed and analyzed using an SRB-assay. The KI library was screened in single technical replicates, and the experiment was repeated in two biological replicates. Rescreening of selected KIs with SRB assay was performed in duplicate technical replicates and the experiment was repeated in three biological replicates.

### 2.11 Bottom-up proteomics sample preparation

For bottom-up proteomics, peptides were prepared by lysis of cells with a 5% SDS solution and Roche cOmplete™ Mini EDTA-free Protease Inhibitor Cocktail (Merck Darmstadt, Germany), followed by sonication (10 minutes, every 30 seconds) (Bioruptor Pico Diagenode, Belgium). Next, protein lysates were quantified using a modified Pierce Micro BCA assay (Termo Fisher Scientifc Rockford, IL), and 5 µg of proteins were used for peptide digestion. Prior to digestion, proteins were reduced with 20 mM dithiothreitol (Merck Darmstadt, Germany) at 45°C for 30 minutes, alkylated with 40 mM iodoacetamide (Merck Darmstadt, Germany) for 30 minutes at room temperature in the dark and acidified with 2.5% phosphoric acid. Finally, proteins were diluted with 90% methanol/100 mM triethylammonium bicarbonate (TEAB) (Merck Darmstadt, Germany) for efficient trapping in Micro S-Trap columns (ProtiFI, Farmingdale, NY, USA). Digestion was performed in the S-Trap overnight at 37°C using Trypsin/Lys-C Mix Mass Spec Grade (Promega, Walldorf, Germany), followed by elution in 50 mM TEAB, 0.2% formic acid (FA) and 50% acetonitrile (ACN) (Merck Darmstadt, Germany). Eluted peptides were dry-evaporated and resuspended in 10% FA solution for subsequent tandem mass spectrometry analysis (LC-MS/MS), as described in Supplementary Methods. The MS proteomics data have been deposited to the ProteomeXchange Consortium via PRIDE (accession number PXD040930).

### 2.12 RNA-seq

Total RNA was extracted from cells using the MiRVAna kit (Ambion, Thermo Fisher Scientific). Library preparation was performed using the Illumina TruSeq Stranded total RNA Library Prep gold Kit (20020598, Illumina Inc., San Diego, USA) and Agencount AMPure XP beads (Beckman Coulter, Brea, USA). Library concentration was determined using a Qubit dsDNA BR kit (Thermo Scientific), and the size distribution was examined with an Agilent Bioanalyzer. Libraries were paired-end sequenced (2×75 bp) on a NextSeq500 (Illumina). BclToFastq was used for the preprocessing of the raw data (trimming and filtering), then FASTQ files were checked for read quality and adapters were removed with Trimmomatic. The resulting reads were then mapped to the human reference genome (GRCh38) with STAR mapping tool (version 2.5.3a) and gene counts extracted with HTSeq. Raw RNA-sequencing data have been deposited on GEO database under accession number GSE228106.

### 2.13 Differential expression analysis

Differential expression analysis was performed with R package Deseq2 (version 1.22.2) for RNA-seq data and R package Limma (version 3.38.3) for proteomics data. In all datasets, black and white cases were allowed retaining the 0 for both parental and resistant cells. For RNA-seq data, only genes having a total sample count >10 were retained. Volcano plots were generated using R package *EnhancedVolcano*, principal component analysis and sample correlation analyses were performed with *plotPCA* function of *DeSeq2* R package and *pheatmap* R package (version 1.0.12). Finally, the R-package *ggvenn* was used to count the genes significantly upregulated in common among the cell lines and between RNA-seq and proteomics datasets.

### 2.14 Statistical analysis

Experiments were performed at least 3 times and data are expressed as mean ± SD of 3 experiments performed in triplicate is shown, unless otherwise specified. To compare between two groups a two-tailed unpaired Student’s t-test was used. For multiple group comparisons a ordinary one-way ANOVA multiple comparison test with Dunnet’s post-hoc test was used, unless otherwise specified in figure legends. Statistical significance was set at p <0.05 and is indicated by *, p < 0.05; **, p < 0.01; ***, p < 0.001; ****, p < 0.0001.

## 3. RESULTS

### 3.1 Establishment of PDAC resistant cell lines

To study resistance to paclitaxel in PDAC, resistant cell lines were established by adaptation to a stepwise increase in the exposure dose over the course of 6-12 months (Fig. 1A). For this purpose, Patu-T, Suit-2.007, and Suit-2.028 cell models were used, and gemcitabine was used as an alternative chemotherapy in parallel to paclitaxel. The IC_50_ values, as extrapolated from the dose-response curves for paclitaxel or gemcitabine, ranged from 2 to 16 nM for all parental cell lines (Fig. 1B and Table S2). IC_50_ values for the resistant derivatives GR and PR were in the µM range (1.6-3.0 µM for PR; 0.7-12 µM for GR), with resistance factors >100-fold that remained stable for at least 2 months of culturing in absence of the drug (Fig. 1B and Table S2). We did not observe cross-resistance: GR cells showed similar or even greater sensitivity to paclitaxel as compared to parental cells and PR cells showed similar or even greater sensitivity to gemcitabine as compared to parental cells (Fig. 1C and Table S2). All together these results indicated that resistant models were stable and did not show cross-resistance, therefore providing a valid model to investigate drug-specific chemoresistance mechanisms.

**Figure 1.**
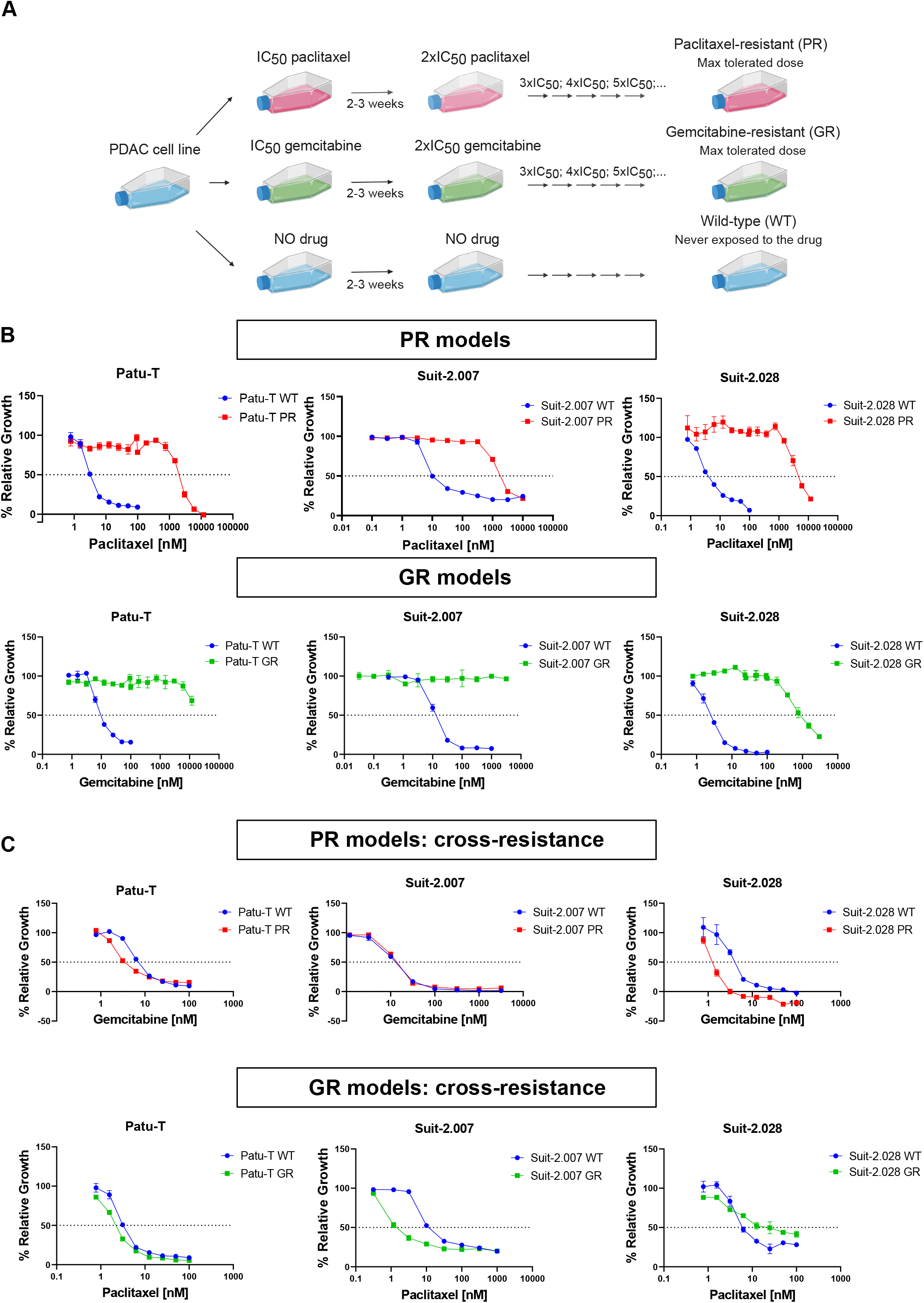
Establishment of paclitaxel and gemcitabine resistant PDAC models. **A**. Graphical representation of the methodology used for generation of PR and GR models. **B**. Growth curves of PR (upper panel) and GR (lower panel) cells, together with the parental cells, exposed to increasing concentrations of gemcitabine or paclitaxel. **C**. Sensitivity of PR cells to gemcitabine and GR cells to paclitaxel. **B-C**. Mean and SD of triplicates are shown. IC_50_ values were calculated as mean of 3 independent experiments, each performed in triplicate.

### 3.2 ABCB1 expression is induced in PR but not GR models

To investigate a common molecular mechanism for paclitaxel resistance in PR cells, Patu-T and Suit-2.28 parental and PR cells were subjected to RNA-seq and proteomics analysis. Both RNA-seq and proteomics were performed in triplicate and correlation plot and principal component analyses showed a good separation among WT and resistant cells (Fig. S1A,B). For differential expression of RNA and proteins, cut off criteria were set at log2FC < -2 or > 2 and p-val < 0.05. RNA-seq analysis identified 720 upregulated genes in PR cells (284 unique for Patu-T; 403 unique for Suit-2.28; 34 in common) (Fig. 2A and Fig. S2). Proteomics analysis identified a total of 5309 (Patu-T) and 5231 (Suit-2.028) unique proteins and 209 proteins were upregulated in PR cells (60 unique for Patu-T; 142 unique for Suit-2.028; 7 in common). Intersection of RNA-seq and proteomics data identified ABCB1 and SRI as the only genes whose expression was upregulated at the transcript and protein level in both PR models (Fig. 2B and Table S3). Other ABC transporters were not upregulated in both cell lines and in both RNA-seq and proteomics data sets (Fig. S3). We validated upregulation of ABCB1 by RT-qPCR and Western Blot. ABCB1 was strongly upregulated at the mRNA and protein level in all PR cell lines, as compared to GR and parental cell lines (Fig. 2C,D). Even though Suit-2.028 GR cells showed some increase in ABCB1 mRNA expression, ABCB1 protein levels were not affected. These findings indicated that induction of ABCB1 is a common event in PR PDAC models but not in GR models.

**Figure 2.**
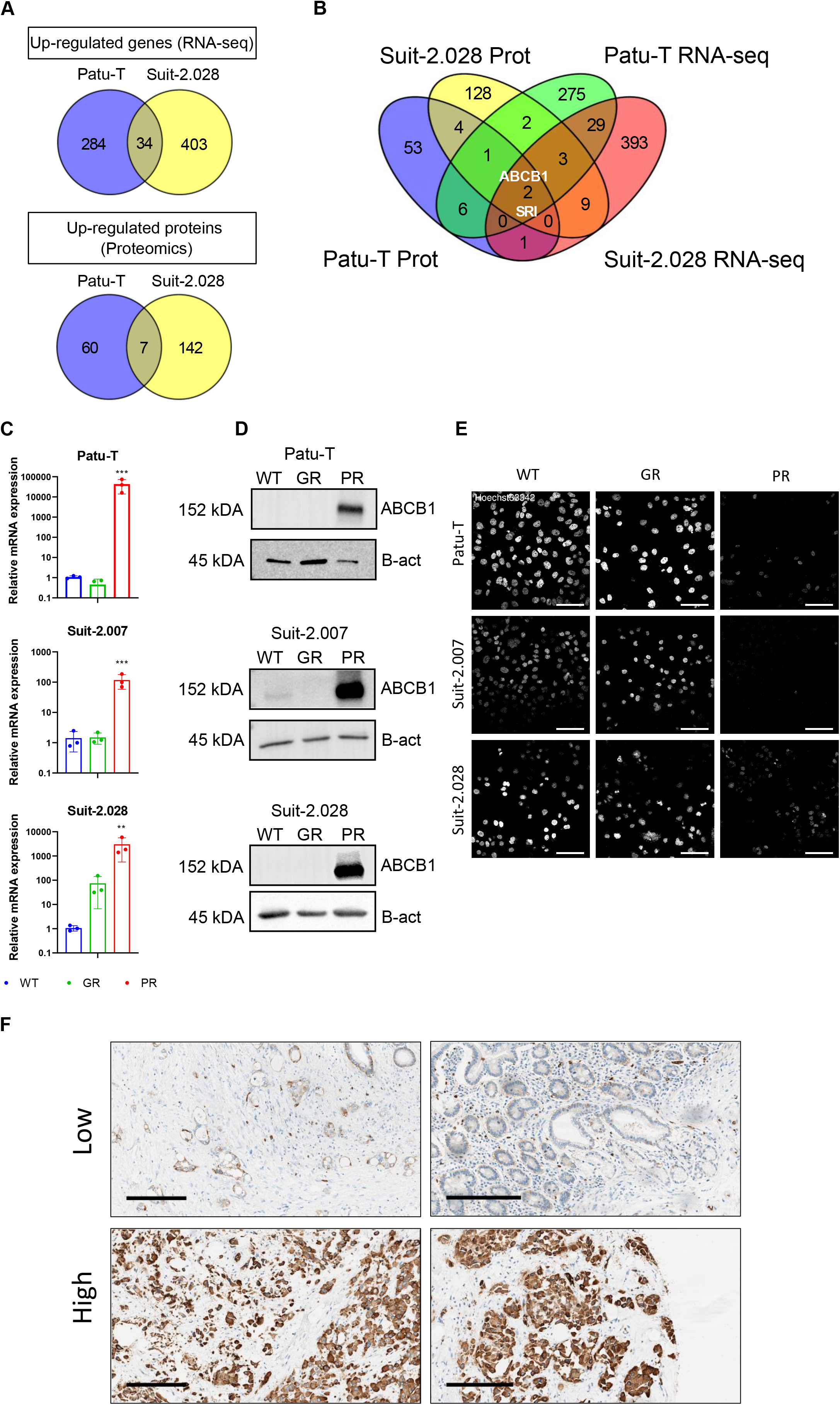
ABCB1 overexpression in PDAC PR cell lines. **A**. Venn diagrams showing the up-regulated mRNAs (upper panel) or proteins (lower panel) in Patu-T PR and Suit-2.028 PR compared to the respective WT. A list of the genes in common for each dataset can be found at Supplemental Table 3. **B**. Venn diagram showing targets upregulated in both RNA-seq (Patu-T PR in green and Suit-2.028 in red) and proteomics analysis (Patu-T PR in blue and Suit-2.028 in yellow) in Patu-T PR and Suit-2.028 PR compared to the respective WT. **C**. Relative gene expression of *ABCB1* in WT (blue), GR (green) and PR (red), measured by RT-qPCR. **D**. Western blot analysis of ABCB1 expression and β-actin as loading control in the indicated WT, GR, and PR cell models. Uncropped Western blot membranes can be found in Figure S5. **E**. Representative confocal images of PDAC cell lines stained with 1 µM of Hoechst33342 for 2h at 37 °C in growth medium. *Scale bar:* 100 µm. **F**. ABCB1 expression levels assessed by immunohistochemistry (IHC) in surgical specimens from PDAC patients treated with gemcitabine/nab-paclitaxel. Two specimens with low and two specimens with high expression levels are shown. *Scale bar*: 200 µm.

### 3.3 ABCB1 represents a target for sensitization of in PDAC to paclitaxel

The functional consequence of increased ABCB1 expression was determined using a Hoechst-efflux assay. The live nuclear stain Hoechst33342 is a substrate of multiple ABC-transporters, including ABCB1 (Shapiro and Ling 1997). Hoechst33342 readily stained nuclei in WT and GR cells but was effectively excluded in PR cells after 2h incubation (Fig. 2E). To confirm the clinical relevance ABCB1, its expression was evaluated in surgical specimens from PDAC patients who received at least one cycle of gemcitabine/nab-paclitaxel therapy. ABCB1 was expressed at different levels in these patients confirming its potential role as a personalized target for therapeutic intervention in PDAC patients (Fig. 2F). There was a trend towards a correlation with poor survival although in this small cohort this was not significant (p = 0.0785, Supplementary Fig. S4A,B). To establish the role of ABCB1 in PDAC paclitaxel-resistance, ABCB1 activity was inhibited with the ABCB1 inhibitor, verapamil (Bellamy 1996).Treatment with verapamil alone did not affect PR cell proliferation (Fig. S6A), but led to a marked increase in Hoechst nuclear staining in PR cells (Fig. 3A,B). In agreement, a dose-dependent increase in the sensitivity to paclitaxel was observed causing a ∼100-1000-fold decrease in the paclitaxel IC_50_ when combined with 10 µM verapamil (Fig. 3C,D). Moreover, verapamil did not affect gemcitabine sensitivity in Patu-T GR or WT (Fig. S6B). As verapamil may have off-target effects in addition to ABCB1 inhibition, the role of ABCB1 in PDAC paclitaxel resistance was further confirmed using gene silencing. Indeed, siRNA mediated ABCB1 depletion strongly sensitized PR cells to paclitaxel as compared to controls (Fig. 3E). Together, these data confirmed the specific role of ABCB1 induction in paclitaxel resistance as a common mechanism underlying paclitaxel-resistance in all three PDAC models.

**Figure 3.**
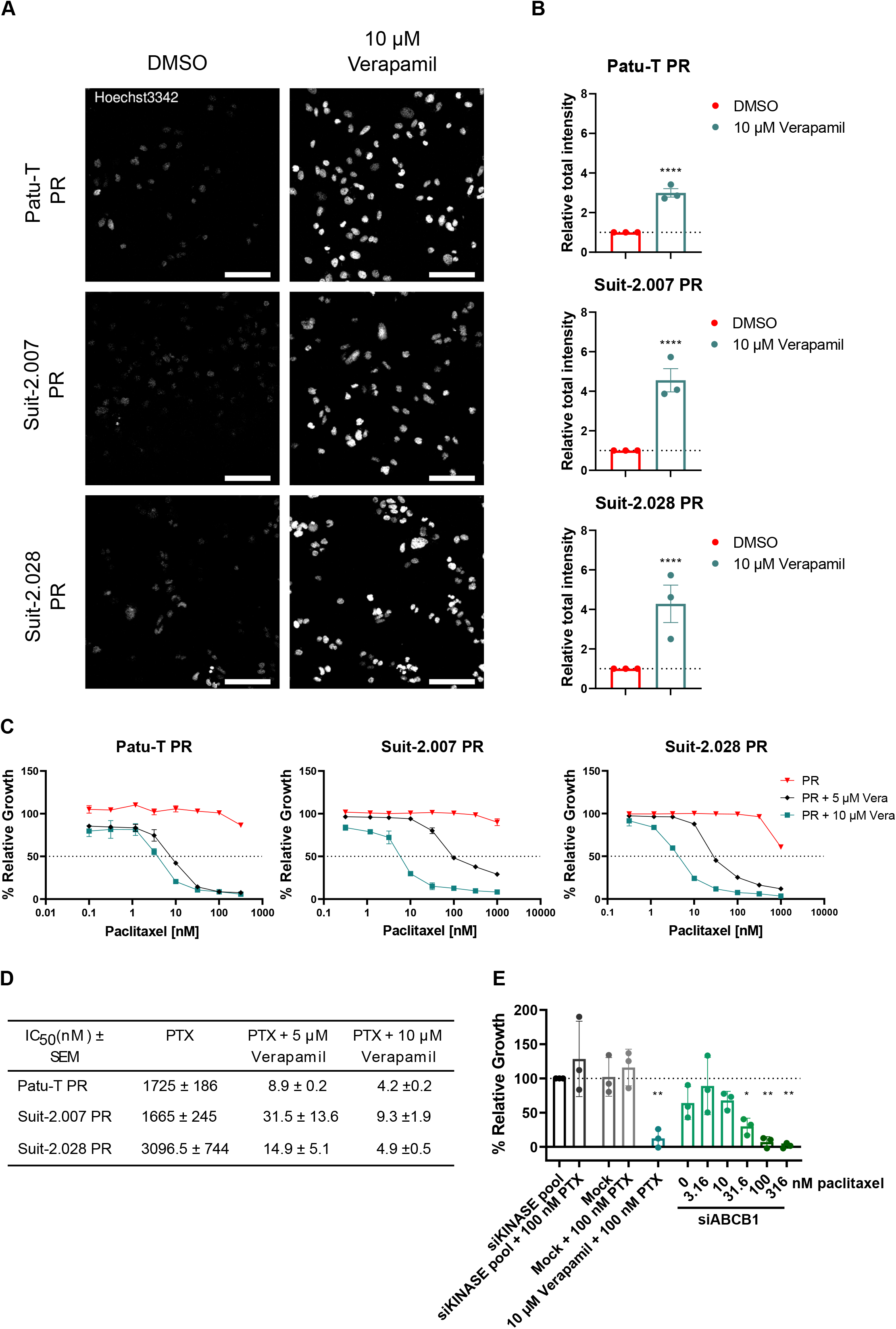
ABCB1 inhibition restores PR cell lines paclitaxel sensitivity. **A**. Representative confocal images of PDAC PR cell lines treated O/N with either verapamil 10 µM or DMSO as a control and stained with 1 µM of Hoechst33342 for 2h at 37 °C in growth medium. *Scale bar:* 100 µm. **B**. Quantification of Hoechst signal total intensity with CellProfiler upon DMSO (red) or 10 µM verapamil (green) treatment. **C**. Representative growth curves of the 3 PR resistant cell models exposed for 72h to paclitaxel concentration ranges, combined with 0.1% DMSO (red triangles), 5 µM verapamil (black diamonds) or 10 µM verapamil (green squares). Mean and SD of triplicates is shown. **D**. Concentrations of paclitaxel causing 50% reduction in cell growth determined in absence or presence of 5 µM or 10 µM verapamil. Mean ± SEM for 3 independent experiments is displayed. As 50% growth inhibition was not fully reached in PR cells exposed to only paclitaxel, values from Supplemental Table 2 are displayed. **E**. Relative proliferation (compared to siKINASEpool control) of Patu-T PR cells 72 hours post-transfection with the indicated siRNA SMARTpools (50 nM) and paclitaxel concentrations, analyzed by SRB assay. Verapamil is used as positive control for ABCB1 inhibition.

### 3.4 ABCB1 gene locus is amplified and gene expression is upregulated in PR cells

ABCB1 overexpression can be caused by amplification of the gene locus 7q21.12 in neuroblastoma, lung, and ovarian cancers (Genovese et al. 2017). To elucidate the mechanism of upregulation in the three PR PDAC models, mRNA expression of the *ABCB4, ADAM22, TP53TG1*, and *SRI* genes that reside in the ABCB1 locus (Fig. 4A), was measured by RT-qPCR. Expression of each of these genes was increased in PR cells as compared to the parental cells for each of the three PDAC models (Fig. 4B-D), suggesting amplification or de-repression of the gene locus. We explored locus amplification by DNA-qPCR in the Patu-T model. Increased signals for all four genes were detected in PR but not in GR cells as compared to WT cells, confirming locus amplification (Fig. 4E). We next investigated the presence of ecDNA that has been associated with increased copies of oncogenes and chemo-resistance in many types of cancer. Similar to parental or GR cells, no ecDNA was present in PR cell metaphase spreads (Fig. 4F). This demonstrated that ABCB1 overexpression in PR PDAC cells was caused by ABCB1 locus amplification which does not involve ecDNA.

**Figure 4.**
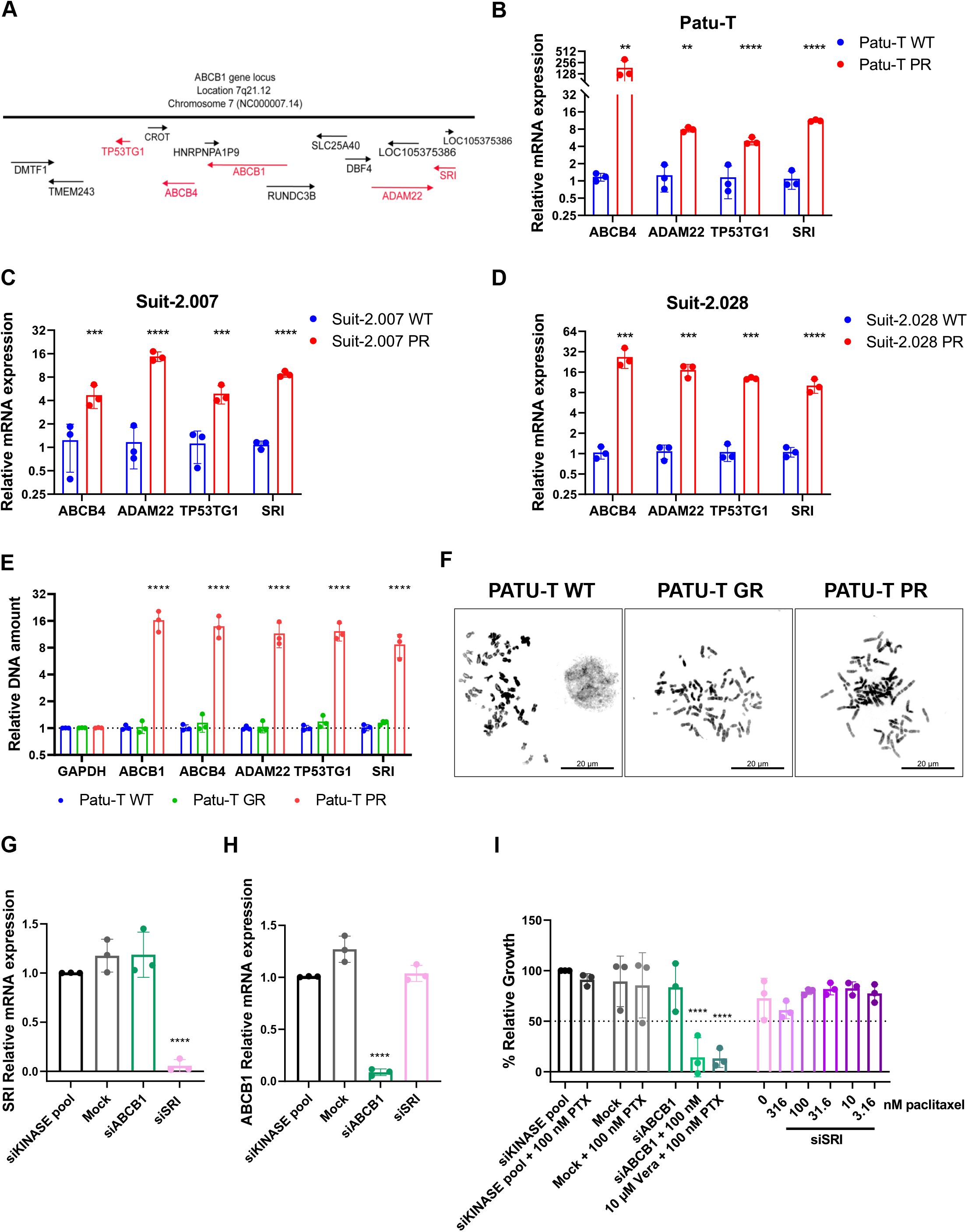
Expression of genes in ABCB1 locus is upregulated in PR cells. **A**. Graphic visualization of ABCB1 amplicon on the locus 7q21.12. **B-D**. Gene expression of *ABCB4, ADAM22, TP53TG1* and *SRI* measured by RT-qPCR in PR (red) relative to WT cells (blue) for the 3 indicated cell models. Brown-Forsythe and Welch ANOVA with Dunnet’s T3 post hoc test was used **E**. Relative DNA amount of *GAPDH, ABCB1, ABCB4, ADAM22, TP53TG1* and *SRI* in Patu-T WT (blue), GR (green) and PR (red) cells, measured by DNA-qPCR and calculated as fold change (2^-ΔΔCt^ compared to the parental). **F**. Representative images of Patu-T WT, GR and PR cell metaphases stained with DAPI. Note absence of ecDNA. *Scale bar:* 20 µm. **G-H**. Gene expression of *SRI* (G) and *ABCB1* (H) in Patu-T PR cells transfected with the indicated siRNA SMARTpools (50 nM) measured by RT-qPCR relative to siKINASEpool samples. **I**. Cell growth of Patu-T PR cells 72 hours post-transfection with the indicated siRNA SMARTpools and paclitaxel concentrations, relative to siKINASEpool control samples. Verapamil (Vera) is used as positive control for ABCB1 inhibition.

### 3.5 Sorcin depletion does not affect proliferation of PR cells treated with paclitaxel

*SRI*, which was the only gene up-regulated at the mRNA and protein level in all three PR PDAC models alongside *ABCB1* (Fig. 2B, Fig. 4B-D), encodes the calcium-binding protein Sorcin that is associated with cancer progression (Battista et al. 2020) and can activate expression of ABCB1 (Yamagishi et al. 2014). We therefore asked if depletion of SRI could reduce ABCB1 levels and restore paclitaxel sensitivity in PR PDAC cells. However, siRNA-mediated silencing of SRI did not lead to reduced ABCB1 expression (Fig. 4G,H) and, in agreement, did not affect paclitaxel resistance of Patu-T PR cells (Fig. 4I). This indicates that a previously described mechanism of ABCB1 regulation by SRI did not underlie ABCB1 mediated paclitaxel resistance in PDAC cells.

### 3.6 Compound screen identifies KIs targeting ABCB1-mediated paclitaxel resistance

Clinical trials with ABCB1 inhibitors on cancer patients have not been successful due to low efficacy or adverse effects (Szakács et al. 2006; Jaramillo et al. 2018). As sorcin targeting proved unsuccessful, we took an unbiased approach to identify alternative pharmacological combinations to restore paclitaxel sensitivity in PR PDAC cells. We screened a library of 760 KIs in Patu-T PR cells. The KI library was screened at a fixed dose of 1 µM in combination with DMSO or paclitaxel at a fixed dose of 0.1 µM. A subset of KIs reduced proliferation of Patu-T PR cells below 50% exclusively when co-administered with paclitaxel (Fig. 5A, Table S4). This subset did not show an enrichment for interaction with specific signaling pathways but several of these KIs had been previously shown to interact with ABCB1. In particular, tyrosine kinase inhibitors, including apatinib and SGI-1776 free base, have been described as ABCB1 inhibitors (Mumenthaler et al. 2009; Mi et al. 2010; Jaramillo et al. 2018; Wu and Fu 2018). We further continued with apatinib and SGI-1776 free base as positive controls and a series of potent KIs detected in the screen for which an interaction with ABCB1 had not been previously shown (Table S4). Sensitization to paclitaxel for PR PDAC cells was validated for each of these selected KIs in the Patu-T, Suit-2.028 and Suit-2.007 models (Fig. 5B and Fig. S7A). In agreement with ABCB1 inhibition by these KIs, each effectively suppressed Hoechst exclusion in the PR cells to a similar extent as that achieved by verapamil (Fig. 5C,D and Fig. S7B). To discriminate between inhibition of ABCB1 efflux function versus inhibition of expression of ABCB1, we measured ABCB1 mRNA expression in the PR models after 48h of treatment with 1 µM of selected KIs. The effect of the KIs varied among cell lines and among KIs, but none of them reduced expression to a level comparable to that in WT cells (Fig. S8). Moreover, changes in expression induced by KIs did not match their effect on Hoechst exclusion or cell proliferation in the presence of paclitaxel, indicating that these KIs primarily act as inhibitors of ABCB1 function. Altogether, these findings identify novel KIs that may be used to target ABCB1-mediated paclitaxel resistance.

**Figure 5.**
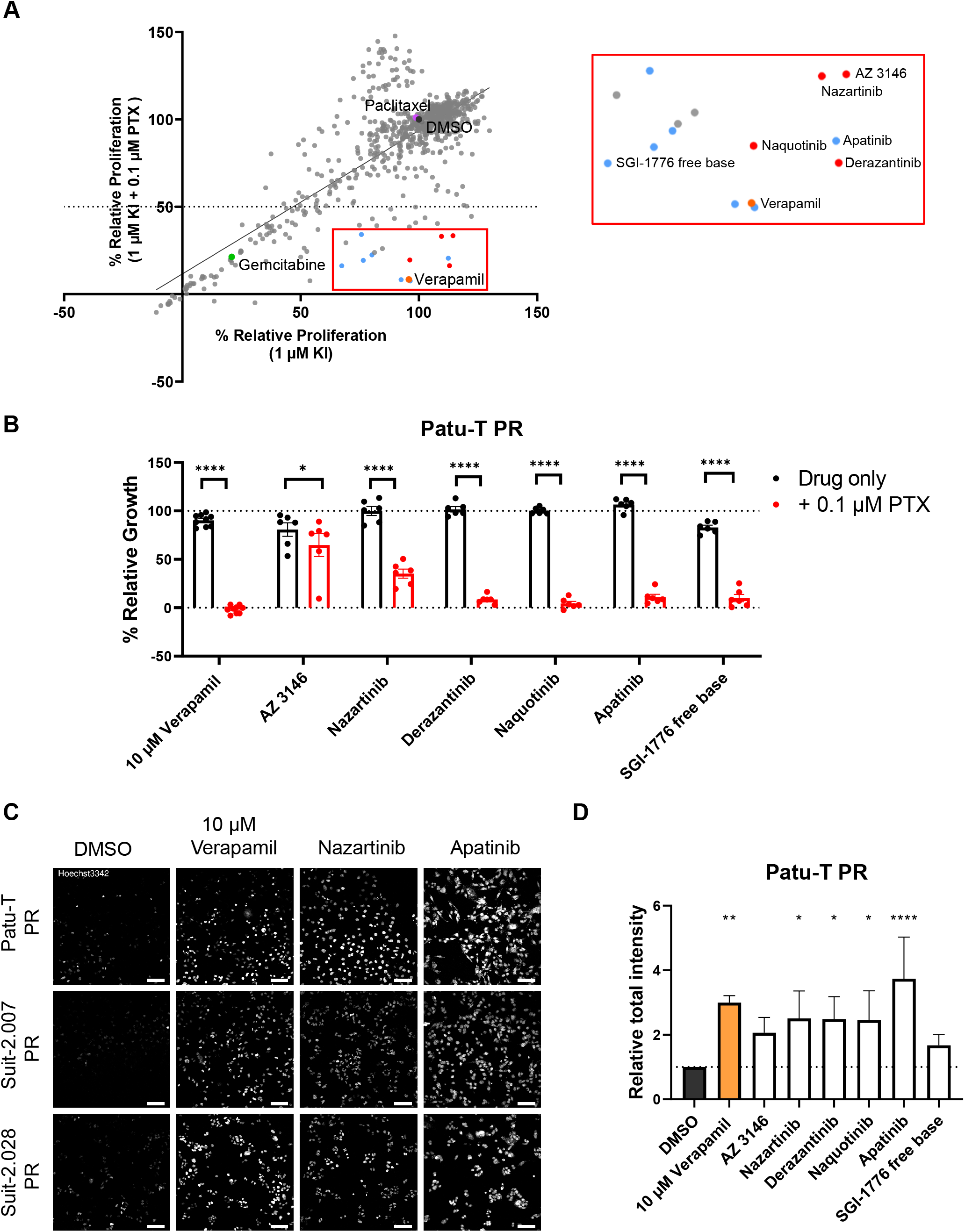
KI library screen to identify synthetic lethalities with paclitaxel in PR PDAC cells. **A**. Scatter plot showing relative proliferation of Patu-T PR cells treated with 760 KIs (1 µM) in absence or presence of 0.1 µM paclitaxel, as assessed by SRB. Dots represent the mean of two independent experiments. Labeled dots indicate 0.1% DMSO control (dark grey), 0.1 µM paclitaxel only (purple), 0.1 µM gemcitabine (green), and 10 µM verapamil (orange). Red box, enlarged on the right, indicates compounds synergizing with paclitaxel, and blue dots indicate compounds already known to interact with ABCB1. **B**. Confirmation of screen hits. KIs were tested at 1 µM concentration in combination with DMSO (black) or 0.1 µM paclitaxel (red) and proliferation was assessed after 72 hours of treatment. Dotted line represents DMSO control (100%). KIs were tested in technical duplicates and controls in triplicates. Mean and SD from 3 independent experiments is displayed (dots indicate individual data points). Ordinary one-way ANOVA was performed, followed by Šídák’s multiple comparisons test. **C**. Representative confocal images of PDAC PR cell lines treated O/N with either 1 µM KIs, 10 µM verapamil or DMSO as a control and stained with 1 µM of Hoechst33342 for 2h at 37 °C in growth medium. *Scale bar:* 100 µm. **D**. Quantification of Hoechst signal total intensity with CellProfiler upon the different treatments. Dotted line represents Relative total intensity = 1.

## 4. DISCUSSION

Chemoresistance is a major hurdle in the treatment of PDAC patients and understanding how to revert resistance to available treatments is of crucial importance. Gemcitabine resistance has been extensively investigated in PDAC, while little research has been performed on paclitaxel resistance (Comandatore et al. 2022; Marin et al. 2022). We find that ABCB1 overexpression is a shared response to continued exposure to paclitaxel in three independent PDAC models and is not associated with gemcitabine resistance in those models. ABCB1 is involved in multidrug resistance of many solid cancers (Gottesman et al. 2002; Vaidyanathan et al. 2016; Sharma et al. 2021). Taxols, in particular paclitaxel, are among the substrates of this transporter. However, the role of ABCB1 in resistance to paclitaxel has not been addressed in pancreatic cancer.

Interestingly, previous studies using pancreatic and other cancer cell lines have shown that ABCB1 is involved in gemcitabine resistance (Chen et al. 2012; Song et al. 2013; Cao et al. 2015; Lu et al. 2019; Marin et al. 2022). Our data do not support such a role: gemcitabine exposure did not induce ABCB1 expression (besides some increase at the mRNA levels in some instances which was not mirrored by enhanced protein levels). Moreover, the induction of ABCB1 in paclitaxel resistant PDAC cells did not lead to cross-resistance to gemcitabine and inhibition of ABCB1 through verapamil did not alter GR cells sensitivity to gemcitabine. The aforementioned studies largely focused on HNF1A or PLK1 mediated gemcitabine resistance mechanisms that involved ABCB1 (Song et al. 2013; Cao et al. 2015; Lu et al. 2019) while Chen et al. did observe increased ABCB1 expression in SW1990 cells treated with gemcitabine (Chen et al. 2012). The contrast between the latter study and our own work is not understood but could be related to the use of different models. Notably, a different study even showed increased sensitivity to gemcitabine in a panel of cancer cell lines overexpressing ABCB1 (Bergman et al. 2003). Altogether, the role of ABCB1 in gemcitabine resistance in PDAC remains unresolved.

Previous studies showed that ABCB1 overexpression can be caused by gene locus amplification (Calcagno and Ambudkar 2010). Indeed, the expression of 4 genes belonging to locus 7q21.12 (*ABCB4, ADAM22, TP53TG1* and *SRI*) are also increased in PR cells. We discriminate between de-repression and amplification by DNA-qPCR, further confirming locus amplification as the underlying mechanism. Gene amplification and chemoresistance have been linked to the presence of ecDNA (Kim et al. 2020). The continuous exposure to a drug like paclitaxel affecting the cell cycle could lead to genomic instability and therefore to the generation of ecDNA fragments (Yan et al. 2020) but we did not find evidence for this. The fact that DNA-qPCR fold-change values were similar for the tested genes in the 7q21.12 locus, while RT-qPCR fold-change values for the same genes differed considerably, suggests that additional mechanisms, on top of locus amplification, may regulate paclitaxel-induced ABCB1 overexpression.

One such mechanism we considered, involves SRI/Sorcin. *SRI* is located on the same gene locus as *ABCB1* and is often co-amplified in multidrug-resistant cancers (Genovese et al. 2017; Battista et al. 2020). In our experiments, SRI was the only candidate specifically induced by paclitaxel along with ABCB1 in the transcriptomics and proteomics datasets for Patu-PR and Suit-2.028 PR. SRI encodes Sorcin, a calcium-binding protein that has been associated with increased tumor aggressiveness (Tong et al. 2015; Battista et al. 2020) and can induce ABCB1 expression in leukemia (Yamagishi et al. 2014). In particular, Sorcin activates Protein Kinase A (PKA)-CREB1 signaling leading to activation of the ABCB1 promoter at cAMP-Response elements (CRE). Our results argue against this mechanism in PDAC paclitaxel resistance: SRI knockdown did not affect ABCB1 expression and failed to sensitize Patu-T PR cells to paclitaxel. It is possible that ABCB1 regulation is different in PDAC cells as compared to leukemic cells or the gradual increase in ABCB1 and SRI caused by paclitaxel differs from engineered SRI overexpression, as used by Yamagishi et colleagues (Yamagishi et al. 2014). Moreover, the impact of sorcin on ABCB1 transcription is modulated by the presence of other mechanisms regulating calcium ion homeostasis, which may vary between cell types (Wang et al. 2022).

Our findings in a small patient cohort show that ABCB1 is expressed in PDAC patients that have received at least one cycle of gemcitabine/nab-paclitaxel treatment. We observe a trend towards a correlation of ABCB1 expression with poor survival, although in this small cohort it was not statistically significant. It will be interesting to assess a larger cohort and compare to patients that have received a different therapy regimen such as FOLFIRINOX. Nevertheless, this indicates ABCB1 is expressed and may represent a target for chemosensitization to paclitaxel in PDAC patients. We confirmed its role as a candidate target for attenuating paclitaxel resistance in PDAC using gene silencing and the ABCB1 inhibitor, verapamil. Unfortunately, verapamil or other ABCB1 inhibitors have not performed well in the clinic. Reasons for the failure of ABCB1 inhibitors in clinical trials include lack of efficacy or dose-limiting toxicity (Szakács et al. 2006; Jaramillo et al. 2018).

Our KI screen identified novel candidate strategies to sensitize PDAC cells to paclitaxel without affecting growth in the absence of the chemotherapy. We found two compounds (i.e., apatinib and SGI-1776 free base) that have already been reported as ABCB1 inhibitors (Mumenthaler et al. 2009; Mi et al. 2010). Interestingly, several compounds that were identified as ABCB1 inhibitors in different cancers failed to sensitize PDAC cells in our screen (i.e., erlotinib (Shi et al. 2007), imatinib (Sims et al. 2013), and nilotinib (Tiwari et al. 2013)). Whether this reflects different inhibitory mechanisms or differences in potency is currently unknown, but it underscores the need to test each drug in the appropriate cancer type. We also identified KIs such as nazartinib, naquotinib, and derazantinib, that have not been previously implicated in PDAC paclitaxel resistance or ABCB1 inhibition. Using a Hoechst efflux assay we confirmed these KIs act by inhibiting ABCB1. For some KIs and in some PR models, inhibition of ABCB1 expression was observed. However, this never reduced it to the nearly absent levels observed in WT cells. Moreover, this effect did not correlate with the efficacy of the KIs in attenuating Hoechst efflux or cell proliferation in the presence of paclitaxel. This indicates that these KIs mainly act by suppressing ABCB1 efflux activity and it points to candidate strategies for combination therapies for chemosensitization. Interestingly, nazartinib, naquotinib, and derazantinib have already passed phase I clinical trials [NCT02108964 (Tan et al. 2020), NCT02500927 (Azuma et al. 2018), NCT03230318 (Mazzaferro et al. 2019)], suggesting that their safety profile is acceptable. Moreover, apatinib is now being tested in phase I and II clinical trials in combination with different chemotherapeutic agents, including paclitaxel [NCT02697838 (Zhao et al. 2020)]. Results from these trials will provide more information on the clinical relevance and feasibility of combining KIs with ABCB1 substrates to improve patient response to therapy.

## Contributors

C.B. designed, executed and analyzed the experiments, prepared figures, and wrote the manuscript; A.G. designed, executed and analyzed the experiments, prepared figures and wrote the manuscript; T.M.S.H. executed experiments; V.vdN., B.vdW., A.Z., I.G. designed experiments; M.C., G.M., A.B., F.F. executed experiments and analyzed -omic results; T.S., L.A.M.D. supervised the study; E.G. designed experiments and supervised the study; E.H.J.D. designed experiments, supervised the study and wrote the manuscript. All authors read and approved the final version of the manuscript.

## Acknowledgements

E.G, E.H.J.D, T.S, A.G. and C.B. were supported by the Dutch Cancer Society (KWF Research Grant # 11957). G.M., and E.G. were supported by Italian Association of Cancer Research (AIRC, IG-grant) and by the Cancer Center Amsterdam Foundation (Proof-of-concept grant). L.A.M.D. and A.B. acknowledge support of the Fondazione Pisana per la Scienza ONLUS, under the project entitled “Precision Exosome Analysis for Early Diagnosis of Pancreatic Adenocarcinoma”. I.G. and E.G. were supported by the European organization for Research and Treatment of Cancer (EORTC GI Group’s Young Investigators grant).

Figure 1 was created with BioRender.com. Imaging data were stored and organized with OMERO Database (Allan et al. 2012).

## Availability of data and materials

RNA-seq data supporting the results of this article is available at Gene Expression Omnibus (GEO) database with accession number GSE228106. The mass spectrometry proteomics data have been deposited to the ProteomeXchange Consortium via PRIDE repository with the dataset identifier PXD040930 and 10.6019/PXD040930. Both datasets are currently private, accessible via token request by the reviewers, and will be publicly available upon publication of the manuscript. All experimental datasets and documents generated in this study are available upon reasonable request to the corresponding author.

