## Supplementary data for "ABCB1 overexpression through locus amplification represents an actionable target to combat paclitaxel resistance in pancreatic cancer cells"

**SUPPLEMENTARY METHODS**

### LC-MS/MS analysis and label-free quantification

Peptide samples were analyzed by nanoLC- MS/MS using an EASY-nLC 1000 interfaced with an Orbitrap Fusion Tribrid mass spectrometer equipped with an EASY spray source (Thermo Fisher Scientific, Waltham, MA, USA). Peptides were loaded onto a PepMap C18 precolumn (2 cm x 75 µm internal diameter (ID), 3 µm particle size, 100 Å pore size) (Thermo Fisher Scientific, Waltham, MA, USA) and separated on a PepMap C18 column (50 cm x 75 µm ID, 2 µm particle size, 100 Å pore size) (Thermo Fisher Scientific, Waltham, MA, USA) heated at 35 °C and with a constant flow rate of 300 nl/min. Peptide separation was carried out using a segmented gradient of 0.1% FA (mobile phase A) and ACN/0.1% FA (mobile phase B) as follows: from 5% to 22% mobile phase B in 104 min, from 22% to 32% in 15 min and from 32% to 90% in 10 min. MS data were acquired in positive ion mode using a spray voltage of 2.0 kV, sheet gas set to 1 to minimize neutral contamination and an ion transfer tube temperature of 275 °C. The MS1 survey scan was acquired using the orbitrap (OT) analyzer within a mass range of 375 – 1200 m/z, resolving power of 120.000 FWHM (at 200 m/z), RF lens value of 60%, maximum injection time of 50 ms and maximum ion count of 400000. MS2 was performed using the TopSpeed method in which the most intense precursor ions (2 – 7 charge states and a minimum intensity threshold of 5000) were isolated with an isolation window of 1.6 *m/z* and fragmented by higher energy collisional dissociation (HCD) at a normalized collision energy (NCE) of 27%. The total cycle time was 3 s. Fragment ion detection was performed in the dual-pressure ion trap (IT) with the maximum number ions set to 2000 and a maximum injection time of 300 ms. A dynamic exclusion of 60 s was enabled to avoid the selection of the same precursor ion during its chromatographic elution.

Raw files were uploaded into Proteome Discoverer software (v2.1) (Thermo Fisher Scientific, Waltham, MA, USA) and queried against the human UniprotKB/Swiss-Prot TrEMBL database (202160 sequences, September 2021) using the SEQUEST database search algorithm. Peptide identification was performed using a mass tolerance of 10 ppm and 0.6 Da for precursor and fragment ions respectively, trypsin/Lys-C as endoproteases and up to two missed cleavages. Cysteine carbamidomethylation was set as a static modification (+57.021464 Da) while methionine oxidation (+15.994915 Da) and protein N-Terminal acetylation (+42.010565 Da) were both set as variable modifications. Peptide spectrum matches (PSMs) were determined using a 1% false discovery rate (FDR), using the Percolator module. Protein abundances were exported from Proteome Discoverer and normalized over the sum within each cell line dataset. The equality of variances was assessed, and selected proteins were tested for significance using a Student’s two-tailed t-test. The MS proteomics data have been deposited to the ProteomeXchange Consortium via PRIDE (accession number PXD040930).

**SUPPLEMENTARY TABLES**

| RT-qPCR | Forward | Reverse |
| --- | --- | --- |
| ABCB1 | GGCTACATGAGAGCGGAGGAC | TTCCGTTGCACCTCTCTGGTC |
| ABCB4 | GAAAGGCCAGACACTAGCCC | ACCATCGAGAAGCACTGTCC |
| TP53TG1 | GAGCTGTCCTAACTCTGCGG | GAGGGTTGGGTACCTTCGTG |
| ADAM22 | CCGCGAAGCACAATGCAG | CAACATGAGTCAACTGCGGG |
| SRI | TCCGCTGTATGGTTACTTTGC | GTGCCAGACATATCTCTATCCAG |
| ACTB | ATTGCCGACAGGATGCAGAA | GCTGATCCACATCTGCTGGAA |
| GAPDH | TCGGAGTCAACGGATTTGGT | TTCCCGTTCTCAGCCTTGAC |
| DNA qPCR | Forward | Reverse |
| ABCB1 | CAAGGCAATTCACAGACACAGG | CACTTCAGTTACCCATCTCGAA |
| ABCB4 | AGCCCAAGGGTTTAGGTACTG | CTAAAGGCTGAGACCGCCAG |
| TP53TG1 | CTCAGATTTTGGTGGCAACTTTTCA | GGAAGCAGCCAACAGCAAATTA |
| ADAM22 | TGAGGGAACCAAAAGCTCCC | CATGGCCCCTCTACCCTACT |
| SRI | TGTTGGGCTCACATGAAGGT | GGATGGGGGTGCCATTCATT |
| ACTB | CACTCCAAGGCCGCTTTACA | CACTCCAAGGCCGCTTTACA |

**Supplemental Table S1.** Primer sequences used for RT-qPCR and DNA-qPCR.

| Cell line |  | IC_50_ Paclitaxel (nM) ± SEM | Paclitaxel resistance factor |  | IC_50_ gemcitabine (nM) ± SEM | Gemcitabine resistance factor |
| --- | --- | --- | --- | --- | --- | --- |
| Patu-T WT |  | 2.23 ± 0.67 | NA |  | 8.33 ± 1.42 | NA |
| Patu-T PR |  | 1725 ± 186 | **774** |  | 2.10 ± 0.16 | - |
| Patu-T GR |  | 0.92 ± 0.58 | - |  | > 12000 | **1440** |
| Suit-2.007 WT |  | 16.46 ± 6.42 | NA |  | 7.09 ± 2.19 | NA |
| Suit-2.007 PR |  | 1665 ± 245 | **101** |  | 9.78 ± 1.40 | - |
| Suit-2.007 GR |  | 0.53 ± 0.50 | - |  | > 3000 | **423** |
| Suit-2.028 WT |  | 5.17 ± 0.46 | NA |  | 3.86 ± 0.28 | NA |
| Suit-2.028 PR |  | 3096.5 ± 744 | **599** |  | 1.65 ± 0.35 | - |
| Suit-2.028 GR |  | 21 ± 5.2 | - |  | 711 ± 116 | **184** |

**Supplemental Table S2.** IC_50_ values of the established resistant cell lines and WT and the respective Resistance factors.

| **Upregulated RNAs in two PR models** | | **Upregulated proteins in two PR models** |
| --- | --- | --- |
| HNRNPA1P9 | DMTF1 | **ABCB1** |
| **ABCB1** | ABCB4 | EEF1A2 |
| AC003991.3 | ADAM22 | FAM83H |
| CROT | MAPK15 | **SRI** |
| CTD-2369P2.8 | TP53TG1_2 | SRXN1 |
| STEAP4 | TP53TG1_1 | TBC1D13 |
| RP11-354M1.2 | FDPSP7 | TMEM120A |
| GRM3 | CTB-167B5.1 |  |
| HOXC13-AS | MDH2 |  |
| AC005522.7 | CACNA2D1 |  |
| AC005076.5 | SEMA3C |  |
| AC034228.4 | POR |  |
| DBF4 | TMEM243 |  |
| KIAA1324L | AC005559.3 |  |
| **SRI** | HLA-DQB1-AS1 |  |
| RP11-66B24.9 | RP11-701H16.4 |  |
| RP11-709A23.2 | AP001610.5 |  |

**Supplemental Table S3.** List of upregulated RNAs and proteins shared between Patu-T PR and Suit-2.028 PR cells. Common hits in RNA-seq and proteomics data are indicated in bold.

| Name | Pathway | Target | Literature connection ABCB1 |
| --- | --- | --- | --- |
| AZ 3146 | Cytoskeletal Signaling | Mps1 | no |
| Nazartinib (EGF816. NVS-816) | Angiogenesis | EGFR | no |
| Derazantinib (ARQ-087) | Protein Tyrosine Kinase | FGFR | no |
| Naquotinib (ASP8273) | Angiogenesis | EGFR | no |
| Apatinib | Protein Tyrosine Kinase | VEGFR.c-RET | Yes (Mi Y. 2010) |
| SGI-1776 free base | JAK/STAT | Pim | Yes (Mumenthaler S. 2010) |

**Supplemental Table S4.** Selected hits from KI-screen.

**SUPPLEMENTARY FIGURES**

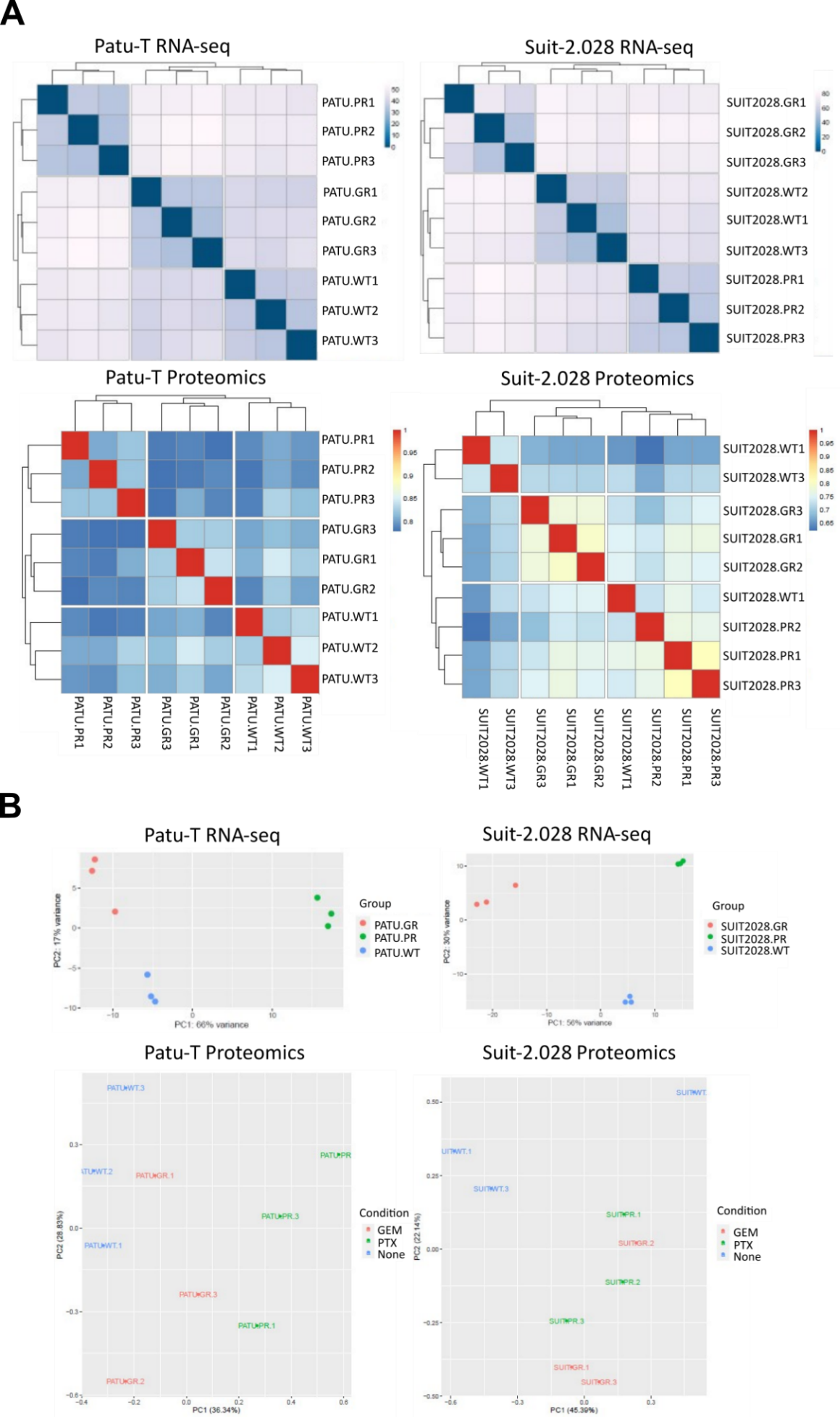

**Supplementary Fig. S1. Correlation plots and Principal component analysis (PCA) of RNA-seq and proteomics data.** **A.** Correlation plots show good correlation for biological replicates within the same cell line, and no or poor correlation among different cell lines. **B.** PCA analysis for RNA-seq and proteomics data. Plots show separation of WT, GR, and PR samples in RNA-seq (top) and proteomics (bottom) data sets.

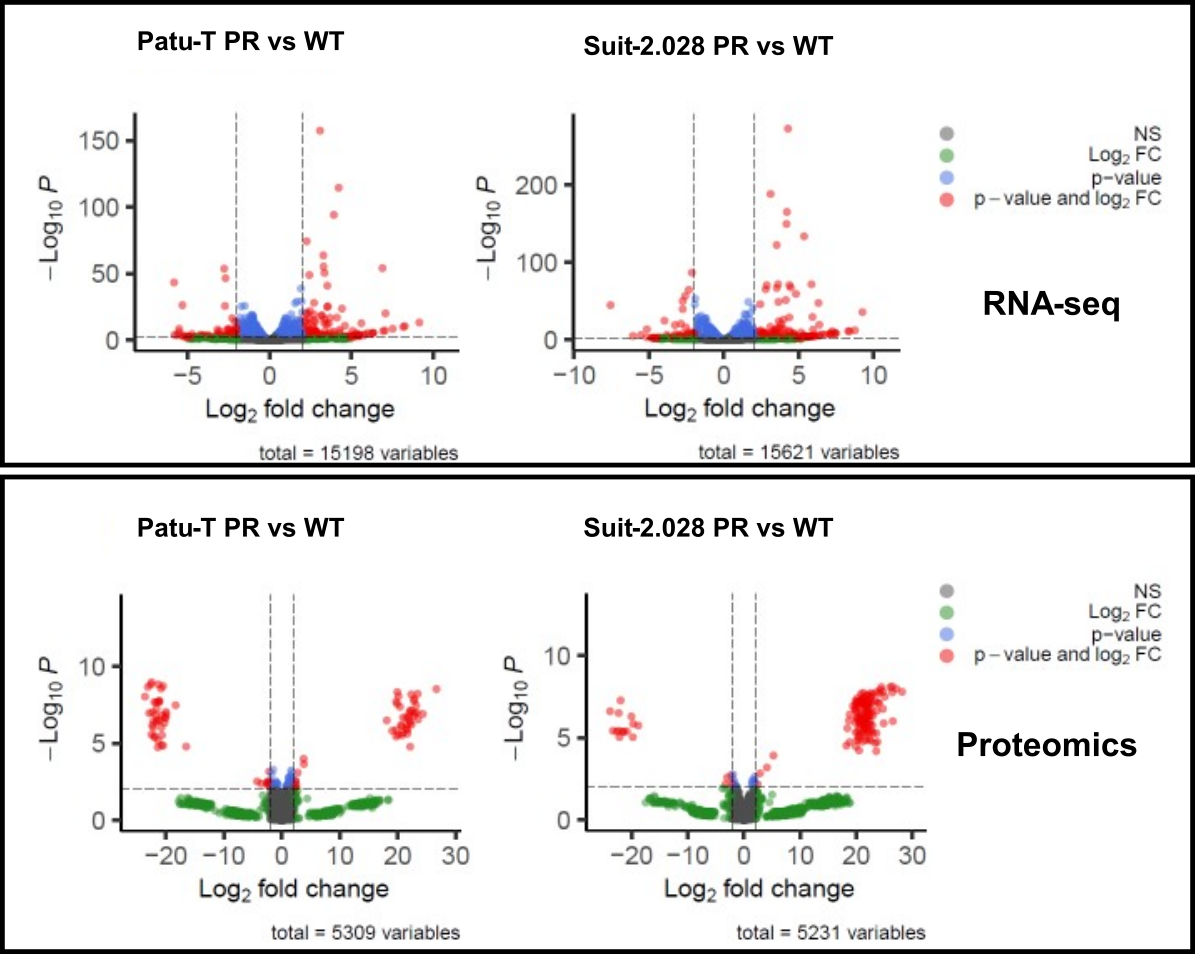

**Supplementary Fig. S2. Volcano plots of differentially expressed genes/proteins for PR vs WT cells.** Grey = not significant (NS); green = only log2FC > 2 or < -2; blue = only p-value < 0.05; red = p-value < 0.05 and log2FC > 2 or < -2. The latter criteria identified 15198 and 15621 differentially expressed RNAs and 5309 and 5231 differentially expressed proteins in Patu-T and Suit-2.028, respectively.

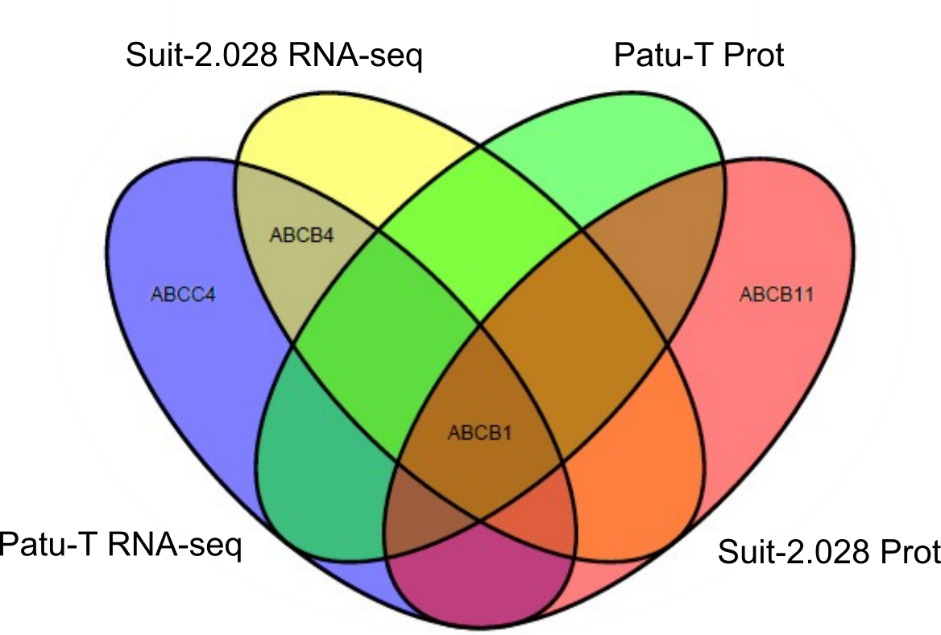

**Supplementary Fig. S3. Upregulation of ABC transporters in PR cells.** Venn diagram showing upregulated ABC transporters in PR models.

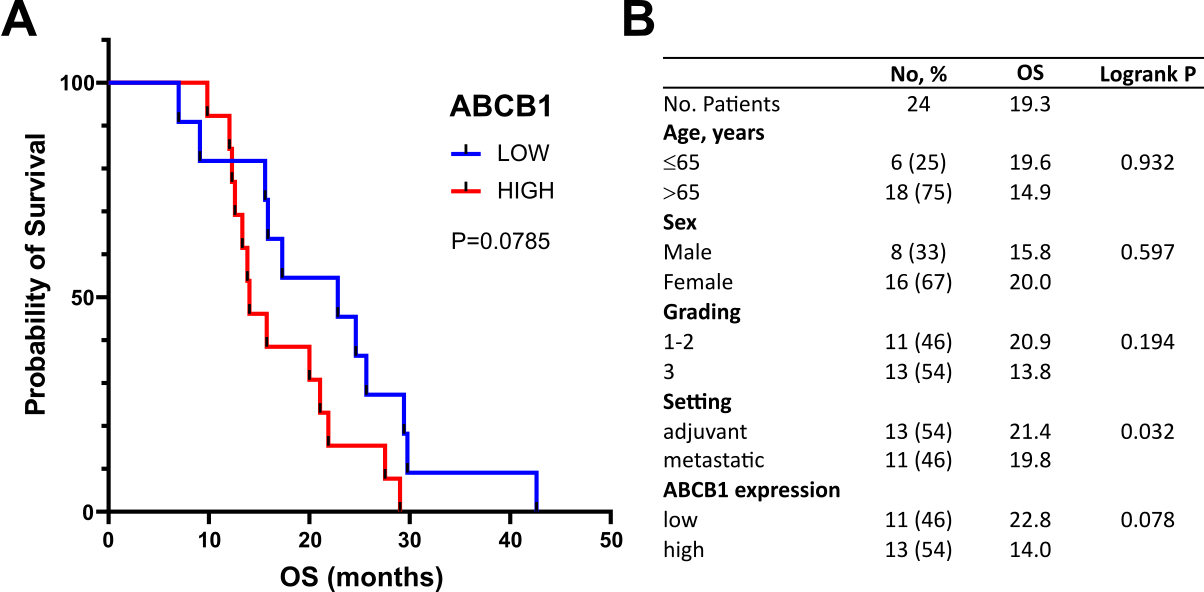

**Supplementary Fig. S4.** **ABCB1 upregulation correlates with poor survival. A.** Kaplan-Meier curves of the patients that underwent surgery, grouped according ABCB1 expression, showing a trend towards reduced probability of survival in case of high (red) vs. low (blue) expression. **B.** Clinicopathological characteristics and correlation with mean overall survival (OS) of the PDAC patients.

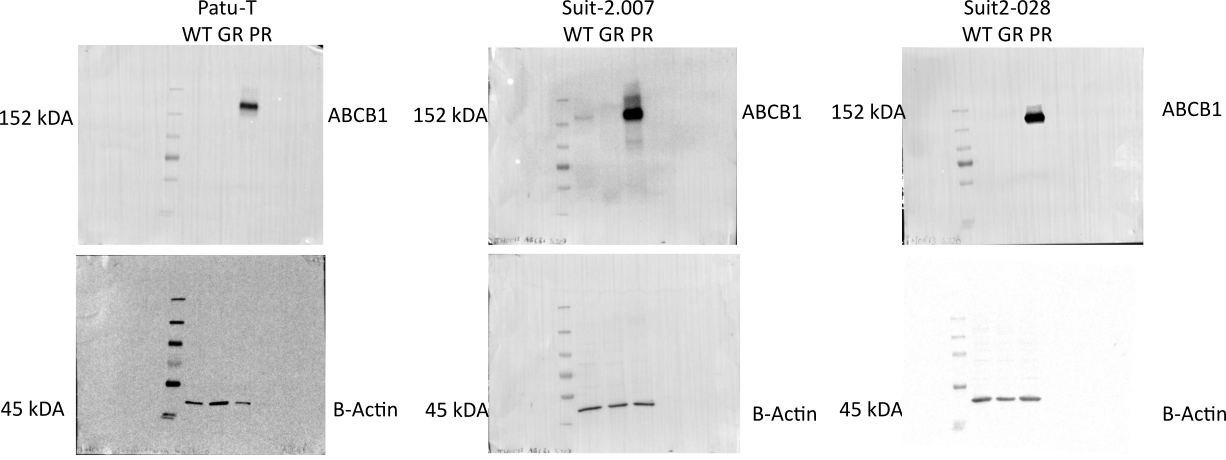

**Supplementary Fig. S5. Uncropped Western blot membranes for Patu-T, Suit-2.028 and Suit-2.007 WT, PR, and GR cells stained for ABCB1 and B-actin.** One biological replicate of western blot is shown for each cell line. Each sample was collected from untreated cells.

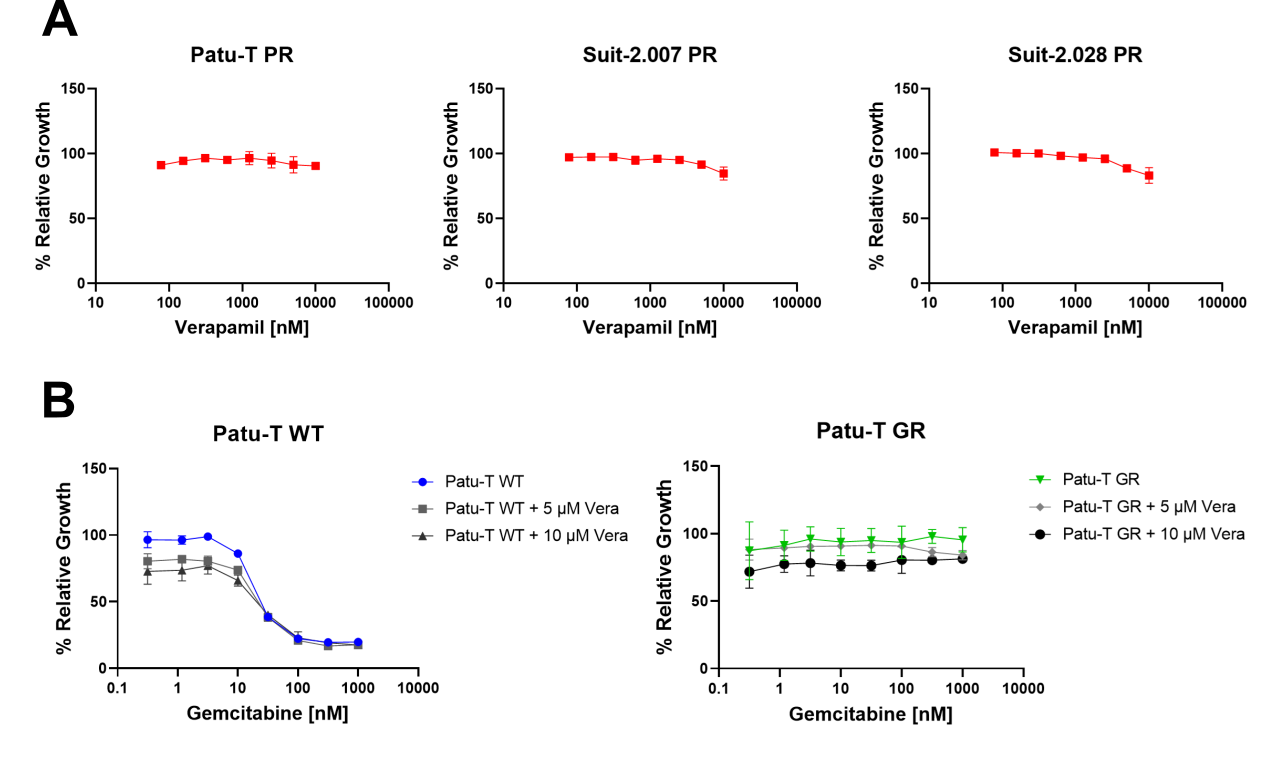

**Supplementary Fig. S6. Verapamil mono treatment does not affect PR cell proliferation** **and verapamil does not sensitize to gemcitabine.** **A.** Representative growth curves of 3 PR cell lines (red) exposed to verapamil concentration ranges, relative to DMSO control. Mean and SD of triplicates is shown. The experiment was repeated 3 times. **B.** Representative growth curves showing effect of 5 µM or 10 µM Verapamil on sensitivity to increasing concentrations of gemcitabine for WT (left) or GR (right) PDAC cells. Mean and SD of triplicates is shown. The experiment was repeated 2 times.

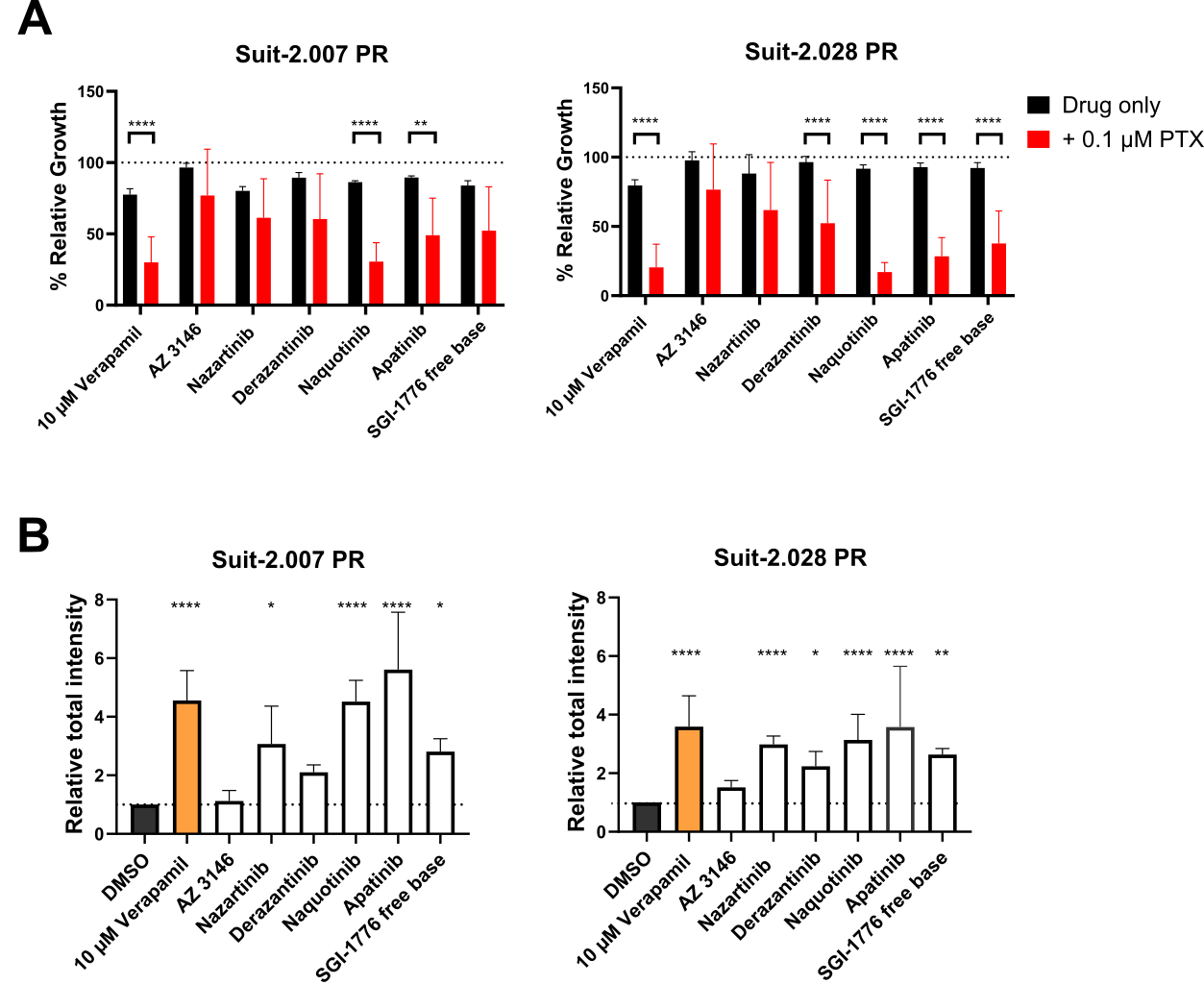

**Supplementary Fig. S7. KI screen validation in Suit-2.007 and Suit-2.028 PR cells. A.** Impact on cell proliferation. Selected KIs from KI library screen were tested at 1 µM in absence (black bars) or presence (red bars) of 0.1 µM paclitaxel (PTX). Proliferation was assessed after 72 hours of treatment. Dotted line represents the DMSO control (100%). Experiments were performed in technical duplicates. Mean and SD of three independent experiments is shown. Ordinary one-way ANOVA was performed, followed by Šídák’s multiple comparisons test. **, p < 0.005; ****, p < 0.0001. **B.** Impact on Hoechst exclusion. Selected KIs from KI library screen were tested for their ability to prevent Hoechst exclusion in PR cells. Relative Hoechst signal intensity for the indicated treatments versus DMSO is shown. Mean and SD of 3 independent experiments performed in triplicates is shown. Ordinary one-way ANOVA with Dunnet’s post hoc test was used. *, p < 0.05; **, p < 0.005; ****, p < 0.0001.

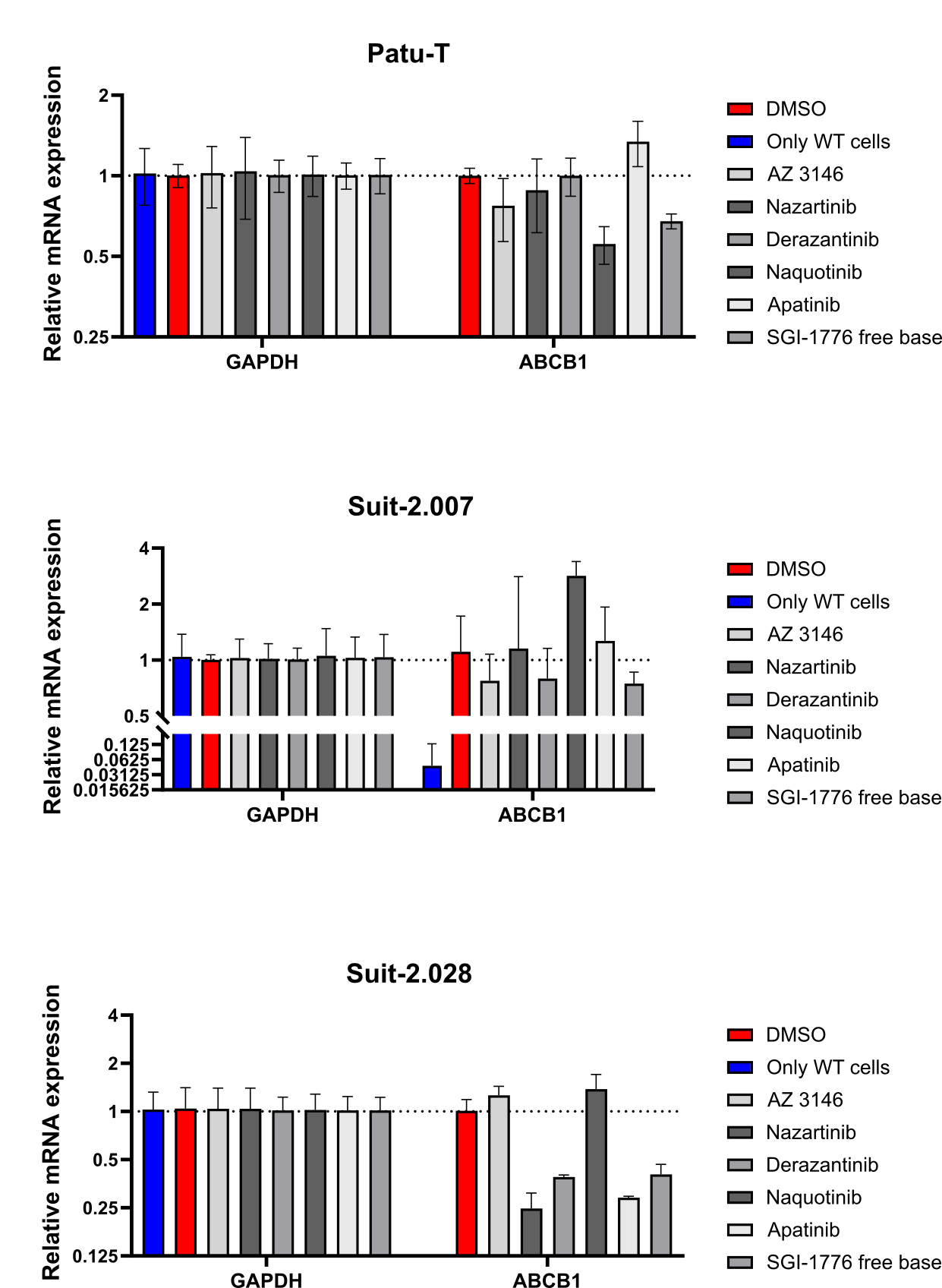

**Supplementary Fig. S8.** KI treatment of PR cells does not decrease ABCB1 expression to level of WT cells. Gene expression of ABCB1 and GAPDH in PR cells after 48h of treatment with 1 µM of the indicated KI, measured by RT-qPCR and calculated as fold change (2^-ΔΔCt^ compared to the DMSO control). Untreated WT sample was included as negative control for ABCB1 expression (blue). *Bars*, mean of triplicates. 1 experiment was performed.
